# Lipoproteins carry endocannabinoids that inhibit the Hedgehog pathway

**DOI:** 10.1101/000570

**Authors:** Helena Khaliullina, Mesut Bilgin, Julio L. Sampaio, Andrej Shevchenko, Suzanne Eaton

## Abstract

Hedgehog proteins are lipid-modified secreted signaling molecules that regulate tissue development and homeostasis. Lipids contained in circulating lipoproteins repress the Hedgehog signaling pathway in the absence of Hedgehog ligand, but the identity of these lipids is unknown. Here, using biochemical fractionation and lipid mass spectrometry, we identify these inhibitory lipids as endocannabinoids. Endocannabinoids are present in lipoproteins of both flies and humans, and repress the pathway in both mammalian signaling assays and *Drosophila* wing imaginal discs. In *Drosophila*, endocannabinoids are required *in vivo* to keep the levels of Smoothened and full-length Cubitus interruptus (Ci_155_) low in the absence of Hedgehog. Furthermore, elevating their endogenous levels inhibits Hedgehog-dependent accumulation of Smoothened and Ci_155_. Interestingly, *cannabis-*derived phytocannabinoids are also potent pathway inhibitors in flies and mammals. These findings constitute a novel link between organismal metabolism and local Hedgehog signaling, and suggest previously unsuspected mechanisms for the broad physiological activities of cannabinoids.

## Introduction

The Hedgehog (Hh) pathway regulates growth, patterning and tissue homeostasis, and its ectopic activation underlies development of a variety of tumors [1,2]. Covalent lipid modifications of Hh ligands are important for their signaling activity [3,4] and allow both *Drosophila* and mammalian Hh proteins to associate with lipoproteins [5]. These particles play multiple conserved roles in regulating Hh signaling. In addition to promoting the release of Hh and its mammalian homologue Sonic hedgehog (Shh), both *Drosophila* and mammalian lipoproteins also contain unknown lipids required to repress the pathway in the absence of lipoprotein-bound Hh ligands [5,6]. The Hh receptor, Patched (Ptc), regulates lipoprotein trafficking in the *Drosophila* wing disc. When Hh is absent, Ptc lowers levels of the 7-pass transmembrane protein Smoothened (Smo) on the basolateral membrane, and blocks Smo-mediated accumulation of full-length transcriptional activator Cubitus interruptus (Ci_155_), by a mechanism that requires lipoprotein lipids [6]. Lipoprotein-associated forms of Hh and Shh block pathway inhibition by lipoprotein-derived lipids [5,6]. Both mammalian and *Drosophila* cells also secrete active, non-sterol-modified forms of Hh (HhN* and ShhN*) independently of lipoproteins. However, these forms cannot block pathway inhibition by lipoproteins [5]. Identifying the endogenous lipoprotein lipids that repress Hh signaling is not only important for understanding the logic of the pathway, but could also suggest new types of pathway inhibitors for clinical applications. Therefore, we have used a combination of biochemical fractionation and lipid mass spectrometry to identify these molecules.

## Results

Since initial experiments showed that very-low density lipoproteins (VLDL) carried lipids that were active in both *Drosophila* and ShhLIGHT2 cells [7] (Figure S1A-B****), we used VLDL as a starting material for fractionation. We exploited both mammalian and *Drosophila* assays to assess pathway inhibition by lipid fractions derived from VLDL. First, we tested their ability to inhibit signaling by non-sterol modified ShhN* in mammalian ShhLIGHT2 cells. Second, we asked whether these lipids could reverse the effects of lipoprotein knock-down in explanted *Drosophila* wing imaginal discs. We showed previously that knock-down of the *Drosophila* lipoprotein Lipophorin (Lpp) elevates Smo levels on the basolateral membrane throughout the anterior compartment of the wing imaginal disc, and does so independently of Hh [6]. Excess Smo signaling then causes ectopic accumulation of full-length transcriptional activator Ci_155_. Both Smo and Ci_155_ accumulation are reversed by addition of protein-free liposomes containing Lpp lipids to explanted wing discs from Lpp RNAi animals (Figure 1A and [6]). Thus, these wing discs represent a powerful *ex vivo* assay for lipids that regulate Smo trafficking and activity.

**Figure 1.**
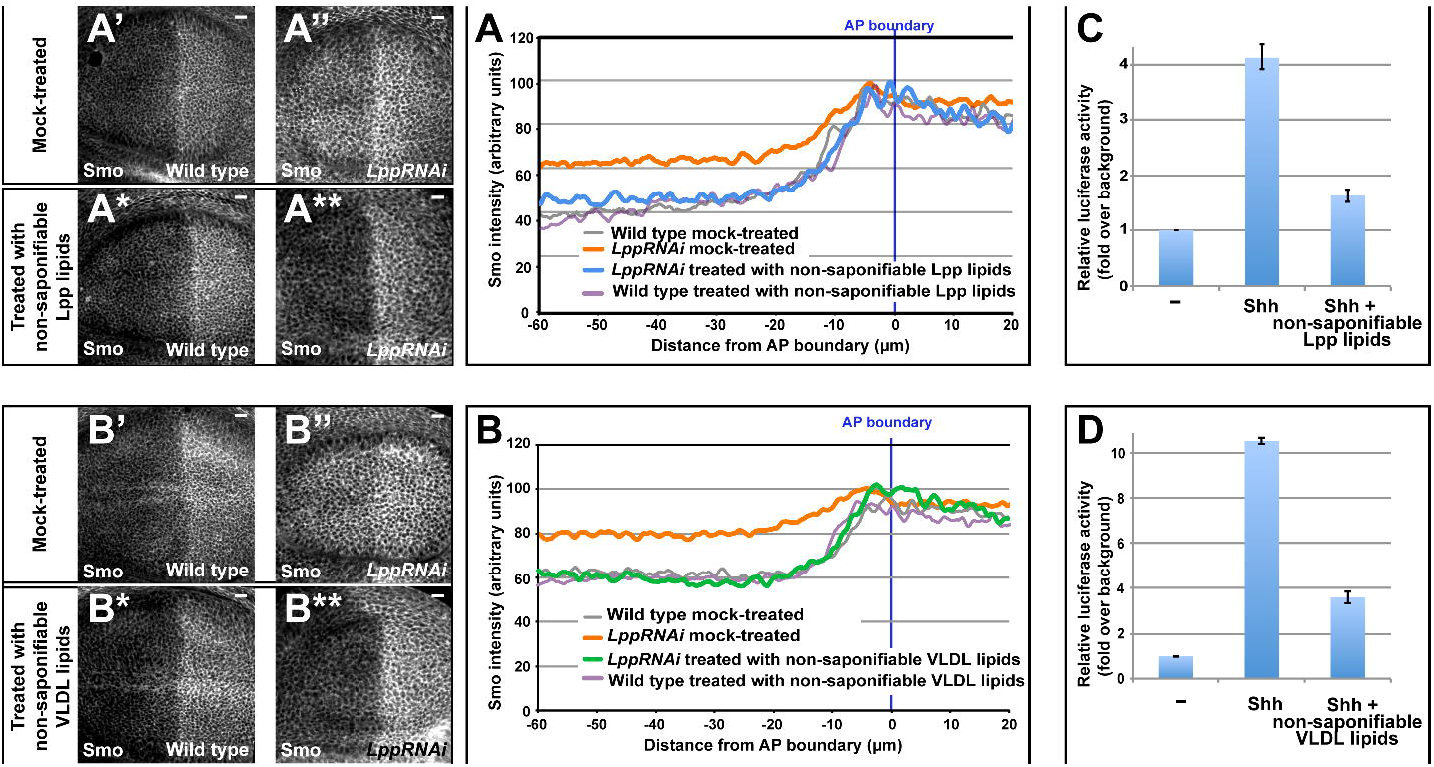
Non-saponifiable VLDL and Lpp lipids reduce Smo levels in *Drosophila* and inhibit Shh signaling in ShhLIGHT2 cells. (A, B) Activities of non-saponifiable Lpp lipids (A, *P* = 1.4×10^−12^) or non-saponifiable VLDL lipids (B, *P* = 4.1×10^−33^) in the wing disc assay. Anterior is to the left, AP: anteroposterior, scale bars = 10 µm. n > 10 discs for each quantification. (C, D) Ratios of reporter activity in ShhLIGHT2 cells treated as indicated. Error bars indicate standard deviations of five independent experiments.

To reduce the complexity of the VLDL lipid pool, we first subjected lipids to saponification and removed the resulting free fatty acids. Saponification depletes glycerolipids containing ester bonds (e.g. tri-, di- and monoacylglycerols, glycerophospholipids and sterol esters), but retains inhibitory activity of lipoprotein lipid extracts in mammalian (Figure 1C and 1D) or *Drosophila* (Figure 1A and 1B) assays. We fractionated saponified VLDL lipid extracts by reversed phase HPLC, assaying elution fractions for their ability to inhibit signaling by non-lipoprotein-associated Shh in ShhLIGHT2 cells. This revealed several clusters of elution fractions with inhibitory activity (red-shaded regions in Figure 2A), and, surprisingly, one region with stimulatory activity (green-shaded region in Figure 2A).

**Figure 2.**
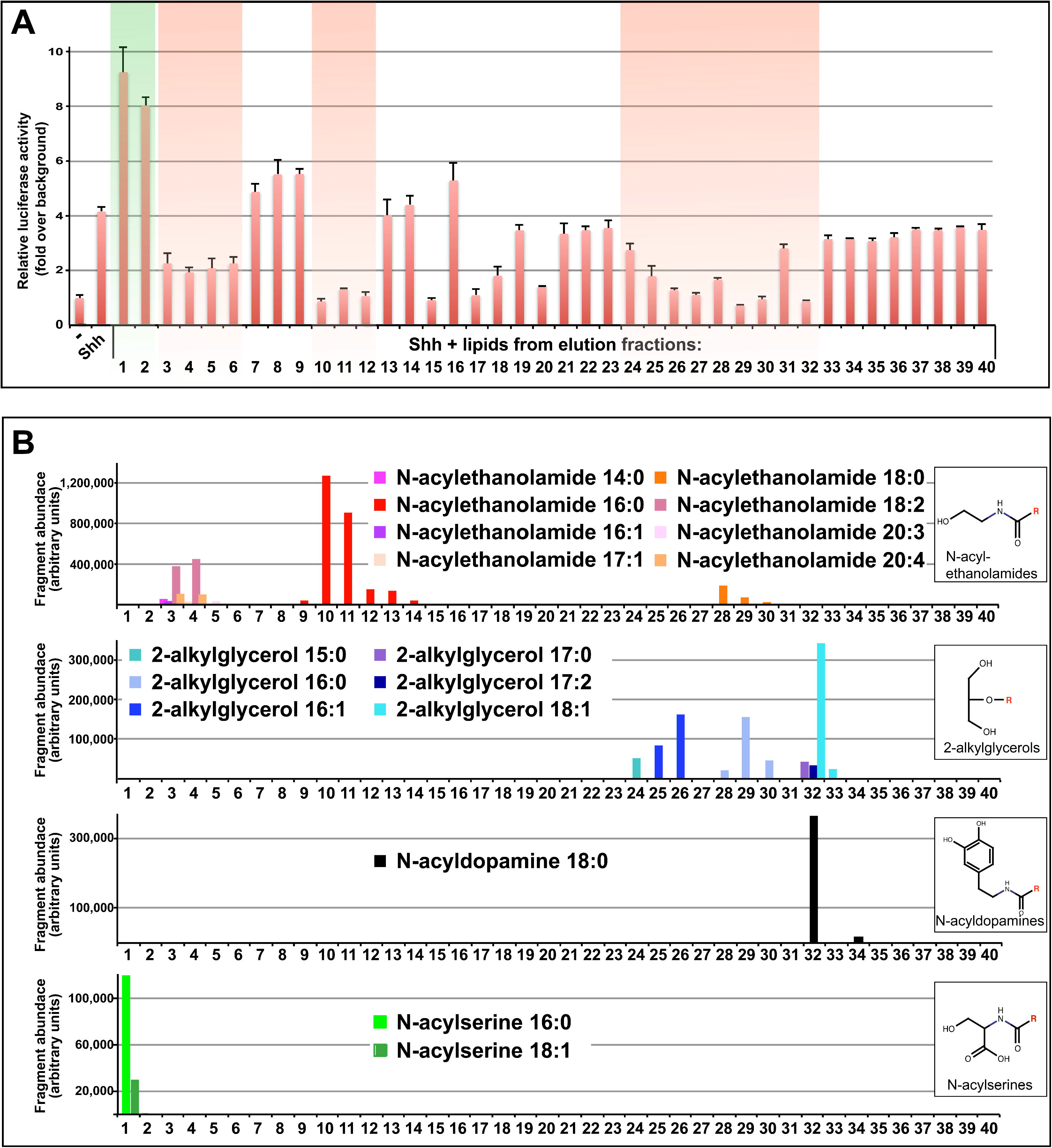
VLDL lipid fractions contain specific endocannabinoids that modulate Shh signaling. (A) Ratio of reporter activity in ShhLIGHT2 cells treated as indicated. Elution fractions with stimulatory/inhibitory activities are shaded in green/ red. Error bars indicate standard deviations of three independent experiments. (B) Distribution of endocannabinoid species, detected by MS/MS in elution fractions tested in (A). Structures of each class are shown in boxes next to distribution profiles. R: species-specific fatty acid/ alcohol moiety.

Since 7-dehydrocholesterol and its derivative Vitamin D3 inhibit mammalian Shh signaling (Figure S1D-F and [8]), we first looked for these compounds using lipid mass spectrometry (MS/MS) and thin layer chromatography (TLC). MS/MS analysis did not detect Vitamin D3 at all. Furthermore, TLC analysis showed that repressive fractions did not coincide with the peak of sterols (not shown). Consistent with this, neither 7-dehydrocholesterol, nor any other sterol, reduced Smo levels in *Drosophila* Lpp RNAi wing discs, and Vitamin D3 had only weak effects (Figure S1G-I). Therefore, Vitamin D3 cannot be the conserved inhibitory lipid we seek.

Hydroxysterols have been shown to stimulate Smo activity [9,10,11] by binding to its N-terminal domain [11,12,13]. Although MS/MS analysis detected hydroxysterols in fractions 33-35, no hydroxysterols were present in stimulatory elution fractions (Figure S2A). Thus, hydroxysterols do not account for the stimulatory activity detected in VLDL and are likely provided by other sources.

Since the active elution fractions did not correspond to peaks of previously reported pathway regulators, we used shotgun analyses by FTMS and MS/MS to search for other signaling lipids (Figure S2G). We detected a variety of sphingolipids, but these did not peak in active fractions (Figure S2B-F) and were inactive in both ShhLIGHT2 and *Drosophila* wing imaginal disc cells (Figure S1J,K). However, MS/MS analysis revealed specific endocannabinoids that co-fractionated with each region of activity (Figure 2 and Figure S2G). Endocannabinoids comprise an expanding family of endogenous compounds that consist of fatty acids and alcohols linked to various polar head groups. Arachidonoyl derivatives of ethanolamine, dopamine and glycerol are potent ligands for the G protein-coupled cannabinoid receptors CB_1_ and CB_2_ [14,15,16]. However, endocannabinoids with different fatty acid moieties and head groups also have interesting biological activities that do not appear to be exerted through cannabinoid receptors [17]. We identified peaks of N-acylethanolamides, N-acyldopamine and 2-alkylglycerols in elution fractions with repressive activity towards the Hh pathway; we also detected peaks of N-acylserines in fractions with stimulatory activity (Figure 2). Thus, VLDL particles carry endocannabinoids that are candidates for Hh pathway inhibitors and activators.

To broadly assess the effects of endocannabinoids on Shh signaling, we assayed synthetically produced endocannabinoids in ShhLIGHT2 cells, including species from each endocannabinoid class found in stimulatory or inhibitory elution fractions. Members of each class potently modulated Hh signaling: N-acylserine 16:0, which we detected in the stimulatory elution fractions, stimulated signaling by ShhN* (Figure 3A). However, it did not activate signaling by itself (not shown). Interestingly, N-acylserine 20:4, which we did not detect in elution fractions, inhibited signaling (Figure S3A). N-acyldopamines, including N-acyldopamine 20:4 (NADA), N-acylethanolamides (including anandamide or AEA), and 2-alkylglycerol 20:4 (noladin ether) repressed signaling by ShhN* (Figures 3B-F and S3B-F). We also tested the activity of 2-acylglycerols (including 2-arachidonoylglycerol or 2-AG), a potent endocannabinoid class that would have been destroyed by saponification of the VLDL lipid extract. 2-acylglycerols also exhibited strong inhibitory activity towards the Hh pathway (Figure S3G, H). Within the N-acylethanolamide and 2-acylglycerol classes, the potency of the inhibition depended on the length/saturation of the fatty acid moiety (Figures 3C-F and S3C-H). Although these compounds were identified on the basis of their ability to inhibit signaling by ShhN*, they also potently inhibit signaling by lipoprotein-associated Shh (Figure S3I and not shown). Taken together, these data indicate that specific endocannabinoids account for inhibitory activities found in VLDL, and that their effects mask the influence of additional, stimulatory endocannabinoids – the N-acylserines.

**Figure 3.**
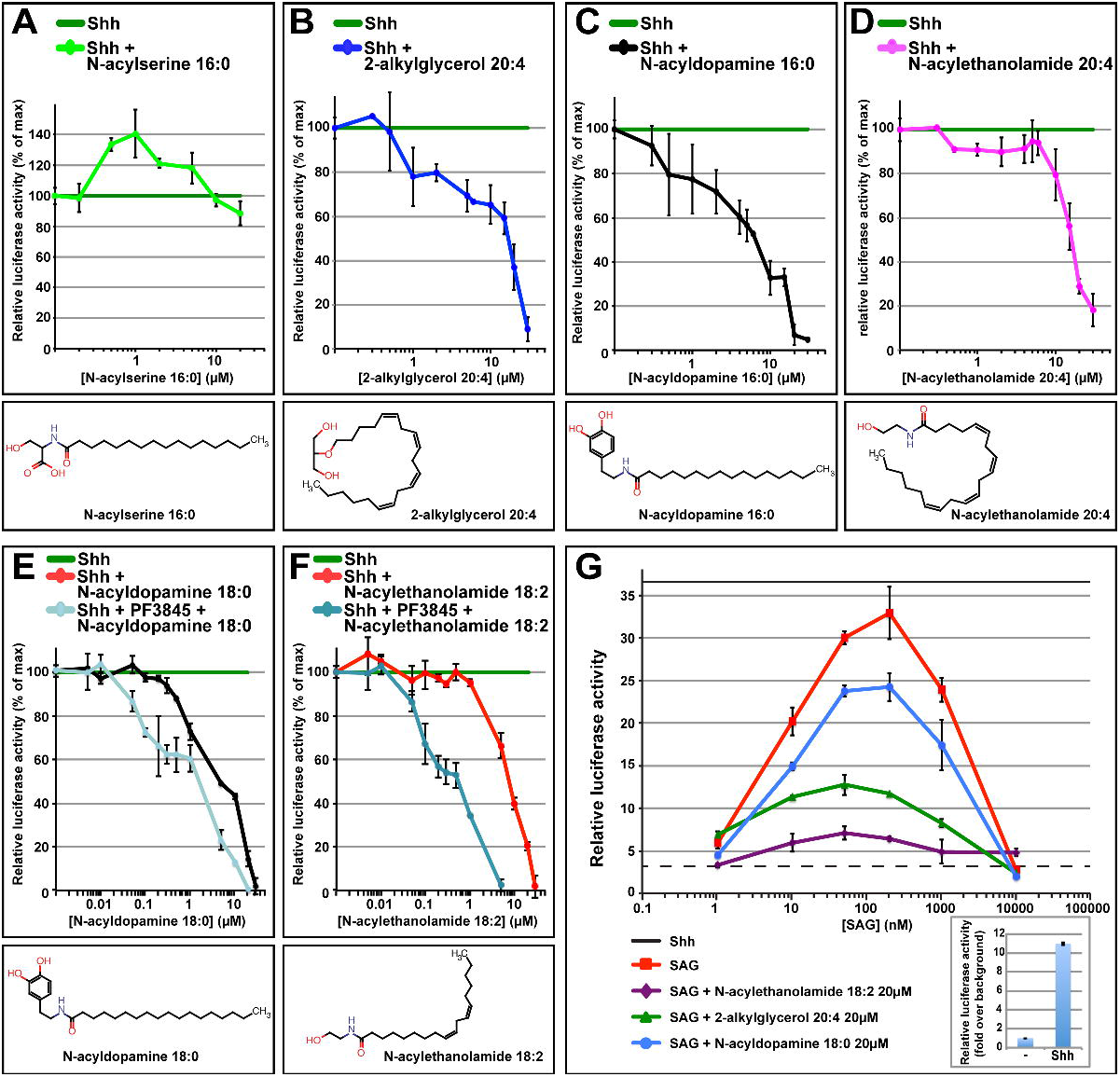
Synthetic endocannabinoids and phytocannabinoids potently modulate Shh signaling and non-competitively inhibit Shh pathway activation by SAG. (A-H) Effects of different cannabinoids and endocannabinoids on signaling by non-lipoprotein-associated Shh in ShhLIGHT2 cells: (A) N-acylserine 16:0, (B) 2-alkylglycerol 20:4, (C) N-acyldopamine 16:0, (D) N-acylethanolamide 20:4, (E) N-acyldopamine 18:0 and (F) N-acylethanolamide 18:2. Blue lines in (E) and (F) show activities of N-acyldopamine 18:0 and N-acylethanolamide 18:2 in the presence of FAAH inhibitor PF-3845. Maximum ratio of reporter activity is normalized to 100 and indicated by green line. In each case, this maximum ratio (i.e. fold stimulation by ShhN* over control) ranged between 8-and 10-fold. Error bars indicate standard deviations of five independent experiments. (G) Hedgehog pathway stimulation by varying concentrations of SAG in the presence of different endocannabinoids. Error bars indicate standard deviations of three independent experiments.

The activity of endocannabinoids on cannabinoid and vanilloid receptors is mimicked by the phytocannabinoids found in *Cannabis.* To examine whether *Cannabis*-derived cannabidiol, cannabinol or delta-9-tetrahydrocannabinol (THC) might also influence Hh signaling, we assayed their effects in ShhLIGHT2 cells. All three compounds potently repressed Shh pathway activity (Figure S3J-L), even in the presence of specific antagonists of CB1 and CB2 (not shown). Taken together, these data suggest that phytocannabinoids may exert some of their physiological effects by repressing Shh signaling.

To study whether phytocannabinoids or endocannabinoids influence the *Drosophila* Hh pathway, we assayed different species of these lipids for their effects on Smo and Ci_155_. Specific N-acylethanolamides and 2-acylglycerols reverse ectopic accumulation of Smo caused by Lpp RNAi, as do cannabidiol and cannabinol (Figures 4 and S3R). These species overlap with those that are active in ShhLIGHT2 cells, but species with slightly shorter fatty acid chain lengths seem to be preferred by *Drosophila* wing disc cells. N-acylserine 16:0, N-acyldopamine 18:0 and 2-alkylglycerol 20:4 (noladin ether) were inactive in *Drosophila* (Figure S3M-P), however we cannot rule out that other fatty acid species might be active. These data show that specific N-acylethanolamide and 2-acylglycerol species repress *Drosophila* Hh signaling by reducing Smo levels and blocking its activity. They further suggest that ectopic accumulation of Smo and Ci_155_ in Lpp RNAi wing discs is caused by impaired Lpp-mediated delivery of endocannabinoids.

**Figure 4.**
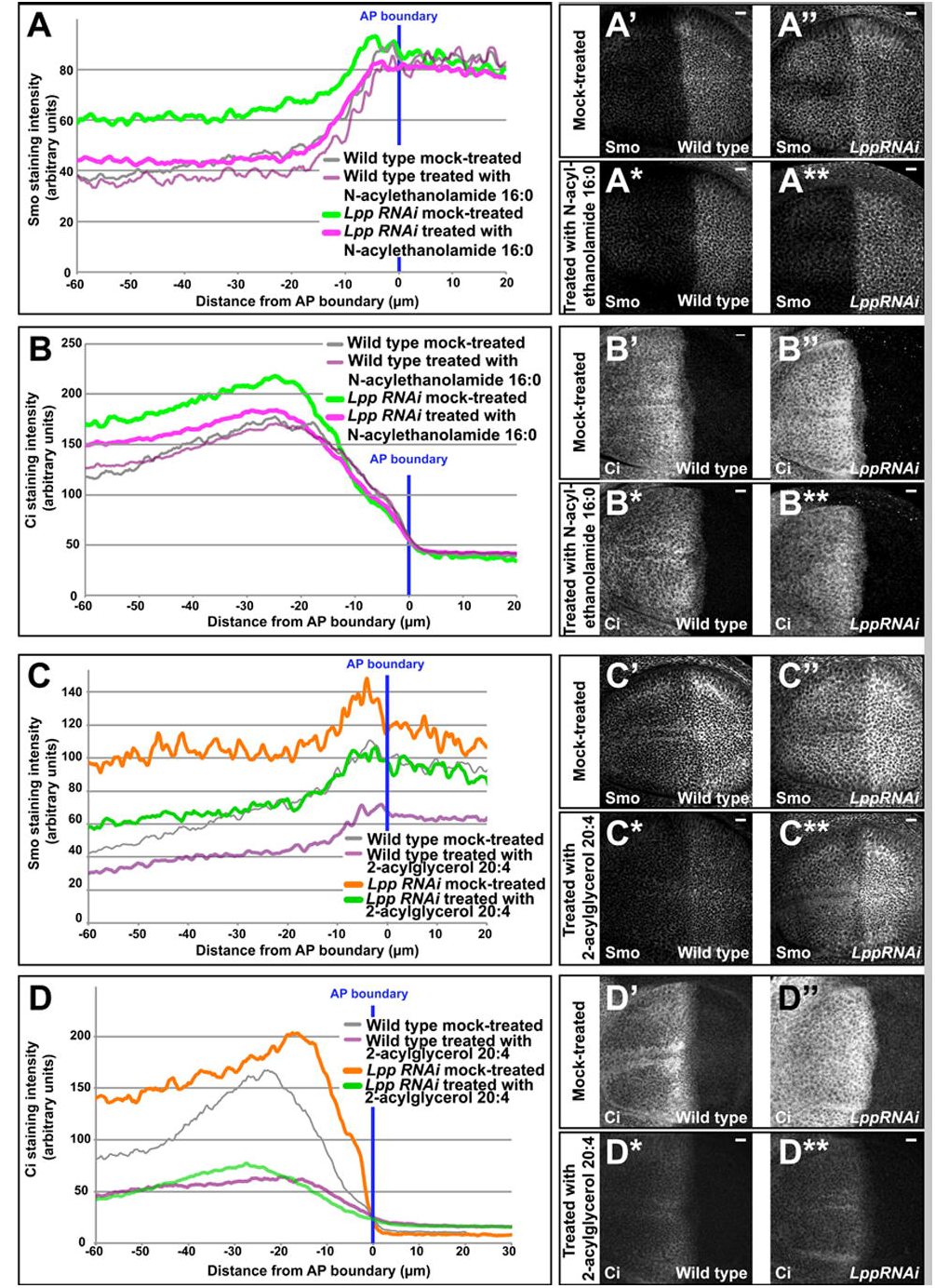
Endocannabinoids present in Lpp reduce Smo and Ci_155_ levels in *Drosophila* wing imaginal discs. (A-B**) Activity of 50 µM N-acylethanolamide 16:0 in the wing disc assay. Anterior is to the right, AP: anteroposterior, scale bars = 10 µm. n > 10 discs for each quantification (*P* = 1.3×10^−37^ in A; *P* = 3.4×10^−2^ in B). (C-D**) Activity of 50 µM 2-acylglycerol 20:4 in the wing disc assay. Anterior is to the left, AP: anteroposterior, scale bars = 10 µm. n > 10 discs for each quantification (*P* = 1.3×10^−18^ for anterior, *P* = 3.8×10^−13^ for posterior in wild type in C; *P* = 2.8×10^−34^ for anterior, *P* = 1.0×10^−20^ for posterior in Lpp RNAi in D).

**Figure 5.**
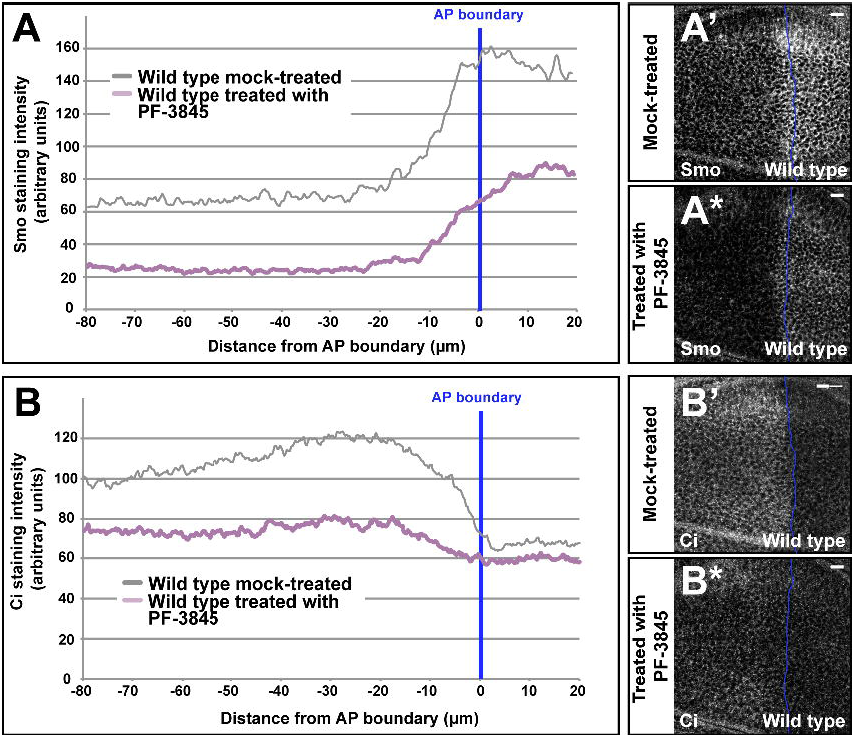
Inhibition of Fatty Acid Amide Hydrolase activity in *Drosophila* wing imaginal discs depletes Smo and Ci_155_. (A-B*) Activity of 10 µM FAAH inhibitor PF-3845 in wild type wing discs. Anterior is to the right, AP: anteroposterior, scale bars = 10 µm. n > 10 discs for each quantification (*P* = 2.56×10^−14^ in A; *P* = 4.82×10^−26^ in B). A/P boundary is marked by blue line.

The most potent endocannabinoids were active at low micromolar concentrations in ShhLIGHT2 cells (Figures 3A-F and S3H). To assess whether endogenous circulating endocannabinoids are sufficiently abundant to suppress Hh signaling *in vivo*, we quantified them in human serum and in *Drosophila* larval hemolymph by LC-MS/MS (Figure S2G-I). As expected, anandamide was present at nanomolar concentrations in human serum [18], however other species containing different fatty acids were much more abundant; total concentrations of N-acylethanolamides, 2-acylglycerols, N-acylserines, 2-alkylglycerols were each in the micromolar range (Figure S2H). Interestingly, the concentration of each endocannabinoid class in the larval hemolymph was also in the micromolar range (Figure S2I). Thus, circulating endocannabinoids are present at levels sufficient to regulate the Hh pathway *in vivo* in both *Drosophila* and mammals.

The concentration at which a ligand is active will depend not only upon its affinity for its receptor, but also on other factors such as access to the receptor and ligand stability. Endocannabinoids are rapidly taken up by cells and metabolized by intracellular enzymes, including fatty acid amide hydrolase (FAAH). This process can deplete even the extracellular pool of endocannabinoids [19,20]. We wondered whether the Hh pathway might respond to lower concentrations of endocannabinoids if their rate of metabolism was reduced. To address this question, we assayed endocannabinoids in the presence of PF-3845, a selective FAAH inhibitor. PF-3845 potentiated the inhibitory activity of endocannabinoids such as N-acylethanolamide 18:2 and N-acyldopamine 18:0, allowing them to repress ShhN*-dependent signaling in the nanomolar range (Figure 3E,F). The low concentration of endocannabinoids required to repress the pathway suggests that these compounds are unlikely to act indirectly by influencing bulk membrane properties. Rather, the active concentrations are consistent with measured binding affinities of different endocannabinoids for receptors such as CB1, CB2, transient receptor potential vanilloid channel (TRPV) and peroxisome proliferator-activated receptor alpha (PPARα) [17]. Therefore, specific ligand-receptor interactions probably mediate the effects of endocannabinoids on Hh signaling.

The *Drosophila* genome encodes 6 proteins with equal similarity to mammalian FAAH, several of which are expressed in wing imaginal discs (Flybase). We wondered whether such enzymes might regulate the metabolism of endocannabinoids in wing discs to tune the Hh pathway. To address this question, we treated explanted wild type wing discs with PF-3845 and monitored the effects on Smo and Ci_155_ 2 hours later. In the anterior compartment, PF-3845 treatment strongly depleted Smo from the basolateral membrane and reduced Ci_155_ accumulation to almost undetectable levels. Interestingly, PF-3845, similar to 2-acylglycerol 20:4 (2-AG), also depletes Smo from the membrane in the posterior compartment, where Ptc is not present. Thus, regulation of endocannabinoid levels by FAAH-like enzymes in the wing disc is crucial for controlling Smo trafficking and Ci_155_ processing. Furthermore, when present at high enough levels, endocannabinoids circumvent the requirement for Ptc.

To investigate how endocannabinoids influence Smo, we asked whether they could prevent pathway activation by Smoothened Agonist (SAG) in ShhLIGHT2 cells. N-acyldopamine 16:0, N-acylethanolamide 18:2 and 2-akylglycerol 20:4 all reduced maximum activation by SAG without affecting its EC_50_. This suggests that endocannabinoids do not compete with SAG for binding to Smo. Rather, they reduce the pool of Smo available for SAG activation.

Endocannabinoids bind to a variety of different receptors including CB1, CB2, TRPV, PPARα and a putative nonCB1/nonCB2 endothelial CB receptor [16,17,21,22,23,24,25,26]. We wondered whether endocannabinoids might act through one of these different pathways to influence Smo. We therefore investigated the effects of specific synthetic agonists and antagonists of these receptors in the ShhLIGHT2 cell assay (Figure S3S). None of these compounds affected either basal or ShhN*-stimulated pathway activity. Thus, endocannabinoids do not act through these receptors to repress the Hedgehog pathway.

## Discussion

Taken together, this work demonstrates that lipoprotein-derived endocannabinoids are conserved endogenous Hedgehog pathway inhibitors that regulate Smo trafficking and block Smo signaling. While oxysterols and Vitamin D have clear effects on the mammalian Hh pathway, endocannabinoids are the first endogenous compounds to show activity towards the Hh pathway that is conserved across phyla.

A long-standing model in the field is that Ptc regulates Smo trafficking and represses Smo activity by controlling the availability of one or more small lipophilic antagonists [2,7,27,28,29]. Thus, one possible explanation for the action of endocannabinoids is that they bind to Smo. This would be consistent with their ability to inhibit SAG-dependent pathway stimulation in ShhLIGHT2 cells. Studies of exogenous and endogenous Smo modulators have demonstrated at least two and possibly three different regulatory binding sites, one of which is utilized by SAG and cyclopamine. If endocannabinoids bind to Smo, they are likely to do so at a different site than SAG, because they reduce peak activation by SAG without altering its EC_50_.

Alternatively, endocannabinoids might act by other more indirect mechanisms to inhibit Smo. Some indirect mechanisms can be ruled out. For example, their ability to block SAG-dependent activation suggests they do not act by stimulating Ptc activity. Furthermore, specific modulators of other known endocannabinoid targets do not affect Shh signaling. Thus, indirect influence via these receptors is unlikely. However, it remains possible that endocannabinoids could act on Smo by specifically regulating as yet unknown proteins involved in Smo trafficking.

Endocannabinoids require chaperones to move through the aqueous environment. There is evidence that serum albumin may carry these molecules [30] and our studies suggest that lipoproteins are also transporters for these lipids. Once present in the plasma membrane, these molecules are rapidly mobilized through the cytoplasm down a concentration gradient by various lipid transfer proteins [31]. This mobilization allows endocannabinoids to be degraded by a number of different enzymes associated with intracellular membrane compartments such as ER, lysosomes and lipid droplets. Intracellular degradation of endocannabinoids is rapid and efficient, and can quickly deplete them from the extracellular environment. It is unclear how endocannabinoids are transferred from albumin or lipoproteins to cell membranes. It is likely that the transfer mechanisms for endocannabinoids in lipoproteins differ from those mediating transfer from other sources, such as albumin. Our previous work has shown that lipoproteins carrying inhibitory lipids (identified as endocannabinoids in this study) are internalized and sequestered in endosomes containing Smo and Ptc. Ptc may act similarly to its relative NPC1 (the protein encoded by the Niemann-Pick type C1 disease gene) – transferring lipoprotein lipids from the endosomal lumen into the endosomal membrane where they can encounter Smo. This sort of transfer mechanism within an endosome might protect lipoprotein-delivered endocannabinoids from rapid transport to compartments where they are degraded.

Competition between different endocannabinoids for these transport and degradation machineries has been shown to give rise to “entourage” effects in which addition of non-ligand molecules can indirectly increase the availability of the real ligand. We have observed that a broad range of different endocannabinoid classes and species influence Smo signaling. It may be that some of the molecules we have identified act indirectly via an entourage effect. The finding that endocannabinoids in systemically circulating lipoproteins repress local Hh signaling demonstrates a fascinating link between systemic metabolism and Hh pathway activity in tissues. It suggests new mechanisms to coordinate organismal growth and development, and also has important implications for adult physiology. The fact that lipoprotein-derived endocannabinoids help to repress such an important tumor-promoting pathway suggests new possibilities to explain the link between disturbed lipoprotein metabolism and cancer risk [32]. In fact, cannabinoids block the growth of many tumors known to depend on Hh signaling [33,34,35,36,37,38,39,40].

Endocannabinoids and phytocannabinoids have a broad range of physiological activities that are not completely understood. Interestingly, Hh signaling regulates many of the same processes -e.g. angiogenesis [41,42], hair follicle development [43,44], nocioception [45,46], bone formation [47,48] and energy metabolism [49,50,51]. Our findings forge a surprising link between cannabinoids and Hedgehog, opening new avenues of research for both important classes of signaling molecules.

## Materials and Methods

### Materials

#### Mammalian cells

ShhLIGHT2 cells [7] were maintained in DMEM + 10% FBS, 150 µg/ml zeocin (Invitrogen) and 400 µg/ml G418 (Invitrogen). ShhLIGHT2 cells are NIH3T3 cells that express Firefly luciferase under the control of a Gli-responsive promoter, and Renilla Luciferase controlled by the viral Thymidine kinase promoter.

#### Mammalian expression plasmids

cDNA encoding human Shh in pCMV-XL5 vector was purchased from OriGene (SC300021).

#### Fly stocks

Oregon R flies, hs-flippase and adh-GAL4 are available from the Bloomington Stock Center. Transgenic line UAS < HcRed > dsLpp is described in [52].

#### Commercially obtained lipids

Sterols: cholesterol, desmosterol, 7-dehydrocholesterol, Vitamin D3 were obtained from Avanti Polar Lipids; ergosterol, ergocalciferol D2, 1,25-dihydroxyvitamin D3 were obtained from Sigma-Aldrich.

All sphingolipids were purchased at Avanti Polar Lipids.

Endocannabinoids: N-acylethanolamides 16:0, 16:1, 18:0, 18:1, 18:2, 20:4, N-acyldopamine 16:0, N-acyldopamine 20:4, 2-alkylglycerol 18:2, 2-acylglycerol 20:4, 2-alkylglycerol 20:4 and N-acylserine 20:4 were purchased at Cayman Chemical, N-acyl dopamine 18:0 was purchased at Tocris Bioscience, N-acyl serine 16:0 was obtained from Avanti Polar Lipids. delta-9-tetrahydrocannabinol was obtained from Biomol Germany. Cannabinol and Cannabidiol were obtained from Sigma-Aldrich. Deuterated endocannabinoids used for LC-MS/MS quantification of endogenous species were purchased from Biomol (Hamburg, Germany) and R&D systems (Wiesbaden, Germany).

All endocannabinoids and Vitamin D3 were stored in dried form at −80 °C and used within 4 weeks after arrival.

ACEA, AM251, AM404, AM630, CB-13, O-1918, CP-775146, GW6471, PF-3845, S 26948 and T 0070907 were obtained from Tocris Bioscience.

Smoothened Agonist (SAG) was obtained from Santa Cruz Biotech.

#### Induction of RNAi

LppRNAi was induced as described [52].

#### Immunohistochemistry

Imaginal discs were fixed and stained as previously described [53]. Antibodies were diluted as follows: anti-Ci 2A1 1:10 [54], anti-Smo 1:50 [55].

#### Image analysis

All quantified immunostaining was performed on discs that were dissected, fixed, stained and imaged in parallel using the same microscope settings. To quantify Ci_155_ and Smo staining intensities, three apical sections 0.7 µm apart were projected using maximal intensity in Fiji. For each image, two rectangles of 100 pixels parallel to the A/P axis by 351 pixels parallel to the D/V axis were selected and centered at the AP boundary in ventral and dorsal compartments. Average pixel intensity was determined as a function of distance from AP boundary using PlotProfile and plotted using Microsoft Excel. All AP boundaries were determined according to anti-Ci_155_ coimmunostaining.

To estimate the significance of changes in staining intensities in discs of different genotypes, we measured Smo or Ci_155_ staining intensity at the same distance from the AP boundary in each disc and calculated *P* values using Excel.

#### Lipoprotein isolation

VLDL, LDL and HDL were isolated from human serum (Sigma) according to [56]. Drosophila Lipophorin (Lpp) was isolated as described previously [6].

#### *Isolation of* Drosophila *larval hemolymph*

Third instar larvae were washed in 10% NaCl and disrupted using a loose dounce homogenizer in 1% acetic acid in H_2_O with 50 µM PF-3845. The loose dounce allows release of the hemolymph but is not designed to disrupt the cells. The extract was centrifuged at 1000 g for 10 min. The supernatant containing hemolymph was centrifuged at 16000 g for 2 h to remove the debris; the supernatant was then subjected to lipid extraction.

#### Lipid extraction

Lipids from purified VLDL, LDL, HDL and Lpp particles applied as crude extracts in ShhLIGHT2 and wing disc signaling assays were extracted by a two-step Bligh and Dyer method [57]. Total lipid concentration was measured according to [58]. Lipoprotein-derived lipids were applied to LIGHT2 cells and wing discs at concentrations found in hemolymph/ human serum.

#### Preparation of saponification-resistant lipids

Dried VLDL lipid extracts were incubated at 80 °C with 2 ml of 0.3 M methanolic potassium hydroxide for one hour. After cooling down to room temperature, the solution was extracted by two washes with diethyl ether. The ether fractions, which contain saponification-resistant lipids, were combined and subjected to further analysis.

#### Drosophila wing disc assay

The assay was performed as previously described [6].

Wing discs dissected from either wild type or LppRNAi animals were incubated with DOPC liposomes (mock) or with liposomes consisting of DOPC:lipids to a ratio of 1:4, at indicated concentration. Non-saponifiable Lpp or VLDL, LDL and HDL lipids were added at concentrations corresponding to those present in the larval hemolymph or human plasma, respectively. After 2 hours at room temperature, discs were fixed and stained for Smo and/or Ci_155_ according to the immunohistochemistry protocol and analyzed as described above.

#### Preparation of Shh applied in the ShhLIGHT2 cell assay

HeLa cells were passaged in DMEM with 10% fetal bovine serum (FBS, Gibco), 50 U/ml penicillin and 50 µg/ml streptomycin (Gibco). To prepare non-lipoprotein-associated Shh, we transfected HeLa cells with plasmids encoding human Shh using polyethylenimine (Polysciences) in OptiMem (Invitrogen), then switched Shh-transfected HeLa cells to serum-free media (DMEM + 1% Insulin-Transferrin-Selenium mixture (ITS-X, Gibco) 4 h post transfection. After 48 h, conditioned media were centrifuged at 1000 g for 20 min and concentrated using Amicon Ultra −10 K (Millipore) for use in the signaling assay. Identically treated media from nontransfected HeLa cells were used as controls. To prepare Lipoprotein-Shh, transfected HeLa cells were cultured in DMEM + 10% FBS. Conditioned media were centrifuged at 1000 g for 20 min. For signaling ShhLIGHT2 assay, lipoprotein-Shh was isolated by density centrifugation [59] and concentrated using Amicon Ultra −10 K.

#### ShhLIGHT2 signaling assay

ShhLIGHT2 cells are NIH3T3 cells that express Firefly luciferase under the control of a Gli-responsive promoter, and Renilla Luciferase controlled by the viral Thymidine kinase promoter. The ratio of Firefly to Renilla Luciferase activity is a specific measure of Hh pathway activity.24 h prior to assay, ShhLIGHT2 cells were plated at 10^5^/ well in 96-well plates and then switched to a serum-free medium consisting of DMEM + 1% ITS-X. In this medium, cells were then treated with nontransfected serum-free HeLa cell supernatants and DMSO or DOPC liposomes (mock) or with non-lipoprotein-associated Shh supplemented with DMSO or DOPC liposomes (Shh) or with different lipids or synthetic compounds.

Purified, commercially obtained lipids or synthetic compounds were added in DMSO at indicated concentration.

Elution fractions were added in DOPC liposomes in a ratio of 1:4.

Luciferase activity was assayed in cell lysates after 24 h, as instructed by the manufacturer (Dual Glo Luciferase Assay, Promega). The resulting Hh pathway activity was measured as the Firefly:Renilla ratio normalized to the ratio in cells that were not stimulated with Shh.

#### SAG competition assay

Smoothened Agonist (SAG) was added to the ShhLIGHT2 cells at indicated concentrations alone or in the presence of different endocannabinoids. Luciferase activity was assayed in cell lysates after 24 h, as instructed by the manufacturer (Dual Glo Luciferase Assay, Promega). The resulting Hh pathway activity was measured as the Firefly:Renilla ratio normalized to the ratio in cells that were not stimulated with Shh.

#### Column chromatography

Extracts containing saponification-resistant lipids from VLDL were loaded onto a C18 column 25 mm x 4.5 mm C18 column packed with 5 µM particles (Vydac, 218TP54). 500 µL of the extract was loaded in 60% aqueous methanol and eluted with a step gradient of 80% methanol in H_2_O (60 min) and a linear gradient of 80% to 100% methanol in water (40 min) at the flow rate of 1 ml/min. The eluate was successively collected into 40 fractions that were dried down and subjected to the ShhLIGHT2 signaling assay.

#### Screening of endocannabinoid-containing fractions by direct infusion mass spectrometry

50 µl of each fraction was mixed with 65 µl of 13 mM ammonium acetate in iso-propanol. Prior to analyses, 20 µl of each sample were loaded into 96-well plate (Eppendorf, Hamburg), sealed with aluminum foil and centrifuged for 5 min at 4000 rpm on a Multifuge 3S-R centrifuge from Heraeus DJB Labcare Ltd (Newport Pagnell, UK). Mass spectrometric analyses were performed on a Q Exactive (Thermo Fisher Scientific, Bremen) instrument; samples were infused by a robotic nanoflow ion source TriVersa NanoMate (Advion BioSciences, Ithaca NY) controlled by the Chipsoft 6.4 software. We used nanoflow chips with a 4.1 µm spraying nozzle diameter; ionization voltage was +/-1.25 kV and gas back pressure was 0.95 psi. FTMS in positive and negative ion mode was acquired with target mass resolution of R_m/z=200_ =140000 within *m/z* range of 100-1000. Precursors within *m/z* range of 200-500 were fragmented in a data-dependent acquisition mode with the target resolution of R_m/z=200_ =17000.

#### Extraction of endocannabinoids from human blood serum and Drosophila larval hemolymph

All operations were performed at 4 °C in the cold room. To 500 µL of human serum or 500 µL of diluted larval hemolymph (an equivalent of the material collected from 13.5 animals) 17.5 µL of the internal standard mixture and 600 µL of ethyl acetate/n-hexane (9:1, V/V) solution were added. The internal standard mixture comprised d4-N-acylethanolamide 16:0 at the concentration of 14 pmol/mL; d8-N-acylethanolamide 20:4 at 7.39 pmol/mL; N-acylserine 20:4 (100 pmol/mL); 2-alkylglycerol 20:4 (100 pmol/mL); d8-2-acylglycerol 20:4 (90.3 pmol/mL) and d5-1-acylglycerol 20:4 (100 pmol/mL. After brief vortexing followed by 15 min centrifugation at 15000 g, samples were incubated on dry ice for 10 min; the upper (organic) phase of two extracts were collected and dried in a vacuum centrifuge. The dried extract was reconstituted in 35 µL of water/ acetonitrile/ iso-propanol/ formic acid (5:4:0.5:0.1, V/V/V/V) mixture, centrifuged for 5 min at 10,000 g and then 30 µL were transferred into a 300 µL glass vial for LC-MS/MS analysis.

#### LC-MS/MS quantification of endogenous endocannabinioids

was performed on an Agilent LC 1100 system (Amstelveen, The Netherlands) interfaced on-line to a triple quadrupole mass spectrometer TSQ Vantage (Thermo Fisher Scientific, Bremen, Germany). We used C18 Zorbax column (5 µm, 0.5 × 150 mm); the flow rate of 20 µL/min; injection volume of 3 µL. The elution gradient was composed from the solvent A: 0.1% formic acid in water; solvent B: acetonitrile/iso-propanol/formic acid (9:1:0.1 V/V/V); the gradient profile was: 0 min, 50% B; 0–2 min, 50-66.4% B; 2–8 min, 66.4-73% B; 8–10 min, 73-95% B; 10-14 min, 95% B; 14-15 min, 95-50% B; 15-21 min, 50% B.

Endocannabinoids were detected as [M + H]^+^ ions by MRM transitions in the positive mode (Table S2 and S3); S-lens voltages and collision energies were optimized individually for each standard compound in direct infusion mode. The transfer capillary temperature was 275 °C and the ion isolation width was 0.7 amu.

MRMs were designed for screening of 26 and 21 endogenous endocannabinoid species in human serum and Drosophila larval hemolymph, respectively, and identified 21 species in serum and 11 species in larval hemolymph (Tables S2 and S3).

## Acknowledgments

We are grateful to Philip Beachy for providing ShhLIGHT2 cells. We thank Barbara Borgonovo for assistance with column chromatography. We thank Petra Born, Natalie Dye, Teymuras Kurzchalia, Andreas Sagner, Wilhelm Palm, Kai Simons, Jonathan Rodenfels and Marino Zerial for critical comments on the manuscript.

## Abbreviations

2-AG: 2-arachidonoylglycerol
AEA: N-acylethanolamide, anandamide
CB: cannabinoid receptor
Ci: Cubitus interruptus
FAAH: fatty acid amide hydrolase
FTMS: Fourier transform mass spectrometry
GPCR: G protein-coupled receptor
Hh: Hedgehog
HPLC: high-performance liquid chromatography
LC-MS/MS: Liquid chromatography-coupled mass spectrometry
Lpp: Lipophorin
MS/MS: tandem mass spectrometry
NPC1: the protein encoded by the Niemann-Pick type C-1 disease gene
PPARα: peroxisome proliferator-activated receptor alpha
Ptc: Patched
SAG: Smoothened Agonist
Shh: Sonic hedgehog
Smo: Smoothened
THC: delta-9-tetrahydrocannabinol
TLC: thin layer chromatography
TRPV: transient receptor potential vanilloid channel
VLDL: very low-density lipoprotein

## Supporting Information

**Figure S1.**

**(A-B****) Lipids derived from mammalian lipoproteins reduce Smo levels in *Drosophila* wing imaginal discs and inhibit Shh signaling in ShhLIGHT2 cells. (A)** Ratio of Firefly:Renilla Luciferase activity in ShhLIGHT2 cells treated as indicated. Treatment with DMSO served as a control for background activity (-). Error bars indicate standard deviations of three independent experiments. (**B**) Quantification of Smo staining in the wing disc assay performed in (**B’-B\*\*\*\***) with lipids derived from HDL (**B\*\*** and magenta in **B**; p = 1.4), LDL (**B\*\*\*** and blue in **B**; p = 0.0048 for far anterior) or VLDL (**B\*\*\*\*** and green in **B**; p = 1.0×10^−^11). Anterior is to the left; AP: anteroposterior, scale bars = 10 µm. n > 10 discs for each quantification.

**(C-I) Sterol addition does not influence Shh signaling in ShhLIGHT2 cells or Smo stability in *Drosophila* wing imaginal discs.** (**C**) Ratio of reporter activity in ShhLIGHT2 cells treated as indicated. Error bars indicate standard deviations of three independent experiments. (**D-F**) Effects of increasing concentrations of 7-dehydrocholesterol (**D**), Vitamin D3 (**E**), and Dihydroxyvitamin D3 (**F**) on signaling by non-lipoprotein-associated Shh in ShhLIGHT2 cells. Ratio of background Firefly:Renilla Luciferase activity (treatment with DMSO alone) is defined as 0. Maximum ratio of reporter activity (ranged between 6-8 fold over background) in cells treated with non-lipoprotein-associated Shh alone is normalized to 100 and is indicated by green line. Error bars indicate standard deviations of at least five independent experiments. (**G**) Quantification of Smo staining in wild type (grey) and Lpp RNAi wing discs that have been mock-treated (orange) or treated with 100 µM cholesterol (red; p = 0.64) 100 µM desmosterol (blue; p = 1.4) or 100 µM ergosterol (green; p = 2.3). Anterior is to the left; AP: anteroposterior. n > 10 discs for each quantification. (**H**) Quantification of Smo staining in wild type (grey) and LppRNAi wing discs that have been mock-treated (orange) or treated with 100 µM 7-dehydrocholesterol (violet; p = 0.57). Anterior is to the left; AP: anteroposterior. n > 10 discs for each quantification. (**I**) Quantification of Smo staining in wild type (grey) and LppRNAi wing discs that have been mock-treated (orange) or treated with 100 µM Vitamin D3 (pink; p = 0.44). Anterior is to the left; AP: anteroposterior. n > 10 discs for each quantification.

**(J, K) Sphingolipids do not influence Shh signaling in ShhLIGHT2 cells or Smo levels in *Drosophila* wing imaginal discs.** (**J**) Ratio of reporter activity in ShhLIGHT2 cells treated as indicated. Error bars indicate standard deviations of three independent experiments. (**K**) Quantification of Smo staining in wing disc assay performed with 50 µM ceramide-phosphatidylethanolamine (red; p = 0.97), 50 µM ceramide 18:1/16:0 (green; p = 0.68), 50 µM ceramide-1-phosphate (violet; p = 0.77), 50 µM sphingosine-1-phosphate (pink; p = 0.83) or 50 µM sphingosine (blue; p = 0.53). Anterior is to the left; AP: anteroposterior. n > 10 discs for each quantification.

**Figure S2.**

**(A-F) Hydroxysterols and sphingolipids do not cofractionate with the inhibitory or stimulatory activities present in VLDL.** (**A-F**) Distribution of hydroxysterols (**A**) and different sphingolipid species (indicated in **B-F**) in elution fractions shown in Figure 2A.

**(G) List of acquisitions and queries used for identification of the different endocannabinoid compounds.** Scripts used for identification and quantification can be found here (https://wiki.mpi-cbg.de/wiki/lipidx/index.php/lx_reference_khaliullina_2012).

**(H)** Endocannabinoids and related molecules in human serum quantified by LC-MS/MS using the method of multiple reaction monitoring (MRM).

**(I)** Endocannabinoids and related molecules quantified in *Drosophila* larval hemolymph by LC-MS/MS by the method of MRM. Concentrations correspond to the values in circulating hemolymph assuming an internal larval volume of 1 µL.

**Figure S3.**

**(A-J) Specific endocannabinoids inhibit signaling in ShhLIGHT2 cells.** Effects of increasing concentrations of (**A**) N-acylserine 20:4, (**B**) N-acyldopamine 20:4, (**C**) N-acylethanolamide 16:0, (**D**) N-acylethanolamide 16:1, (**E**) N-acylethanolamide 18:0, (**F**) N-acylethanolamide 18:1, (**G**) 2-acylglycerol 18:2, (**H**) 2-acylglycerol 20:4 on signaling by non-lipoprotein-associated Shh in ShhLIGHT2 cells. (**I**) Effects of increasing concentrations of N-acylethanolamide 18:2 on signaling by sterol-modified, lipoprotein-associated Shh in ShhLIGHT2 cells. Effects of increasing concentrations of (**J**) delta-9-Tetrahydrocannabinol, (**K**) Cannabidiol and (**L**) Cannabinol on signaling by non-lipoprotein-associated Shh in ShhLIGHT2 cells. Ratio of background Firefly:Renilla Luciferase activity (treatment with DMSO alone) is defined as 0. Maximum ratio of reporter activity in cells treated with non-lipoprotein-associated Shh alone is normalized to 100 and is indicated by green line. In each case, this maximum ratio (i.e. fold stimulation by ShhN* over control) ranged between 8-and 10-fold. Error bars indicate standard deviations of at least five independent experiments.

**(M-P) Some endocannabinoids are inactive in *Drosophila* wing imaginal discs.** (**M-O**) Quantifications of Smo staining in wing disc assays performed with (**M**) 50 µM N-acylethanolamide 18:1 (pink; p = 0.41), (**N**) 50 µM N-acylethanolamide 18:0 (green; p = 0.74) or 50 µM N-acylethanolamide 20:4 (blue; p = 1.6), and (**O**) 50 µM 2-alkylglycerol 20:4 (light blue; p = 2.5). (**P**) Quantification of Smo staining in wing disc assay performed with 50 µM N- acylserine 16:0. Smo levels are unaffected by treatment with N-acylserine 16:0 in both Lpp RNAi (compare orange with green; p = 4.3) and wild type (compare grey with violet; p = 1.9). Anterior is to the left; AP: anteroposterior. n > 10 discs for each quantification.

**(R) Specific endocannabinoids and phytocannabinoids reduce Smo levels in *Drosophila* wing imaginal discs.** Quantification of Smo staining in wing disc assay performed with 50 µM N-acylethanolamide 18:2 (red; p = 1.2×10^−6^), 50 µM 2-acylglycerol 16:0 (brown; p = 1.3×10^−5^), 50 µM Cannabidiol (violet; p = 0.00109) and 50 µM Cannabinol (blue; p = 4.3×10^−14^). Anterior is to the left; AP: anteroposterior. n > 10 discs for each quantification.

(**S**) **Specific agonists and antagonists of non-Smoothened endocannabinoid targets do not influence signaling in ShhLIGHT2 cells.** Signaling by non-lipoprotein-associated Shh in ShhLIGHT2 cells in the presence of specific agonists (green-shaded regions) or antagonists (red-shaded regions) of CB1 and CB2 receptors, nonCB1/CB2 endothelial receptor, PPAR-alpha and PPAR-gamma receptors and TRPV channels. Error bars indicate standard deviations of three independent experiments.

